# Learning and forecasting shared evolutionary pathways to multi-drug resistance across global pathogens

**DOI:** 10.64898/2026.08.30.748110

**Authors:** Olav N. L. Aga, Sabrina J. Moyo, Jakob Fernø, Joel Manyahi, Upendo Kibwana, Iren H. Löhr, Nina Langeland, Bjørn Blomberg, Iain G. Johnston

## Abstract

Infections with bacteria which have evolved multi-drug resistance (MDR) cause millions of deaths worldwide. Large-scale efforts are gathering genotypic and phenotypic data on MDR bacteria, but methods for learning the structure, diversity, and predictors of evolutionary pathways to MDR have yet to take full advantage of these data. Here, we use evolutionary accumulation modelling (EvAM), an emerging class of machine learning methods with roots in cancer progression, to infer these evolutionary pathways across ESKAPEE pathogens (seven bacterial species that dominate health burdens), using a database of over 635k genotyped phenotypic observations from around the world. We identify global patterns in MDR evolutionary pathways, remarkably shared across multiple ESKAPEE species. Species-specific deviations from these stereotypical pathways are connected with geographical and demographic covariates, facilitating predictions of future MDR evolution. We verify these predictions with several hundred new phenotypes from ESKAPEE samples spanning decades of clinical infections in sub-Saharan Africa, demonstrating the capacity to forecast future MDR evolution from these inferred shared pathways.

## Introduction

Drug resistance in the ESKAPEE pathogens (*Enterococcus faecium* [Ef], *Staphylococcus aureus* [Sa], *Klebsiella pneumoniae* [Kp], *Acinetobacter baumannii* [Ab], *Pseudomonas aeruginosa* [Pa], *Enterobacter* sp. [Es], *Escherichia coli* [Ec]) dominates the global health threat from bacterial resistance (Miller & Arias, 2024; Naghavi et al., 2024). The health and economic burdens of multi-drug resistance (MDR, simultaneous resistance to three or more drug families) involves millions of deaths and hundreds of billions of dollars, and is increasing rapidly (Marino et al., 2025). Understanding the evolution of MDR across pathogens, its covariates, and possible future bacterial behaviours, can contribute to efforts combatting this growing global health issue (Baquero et al., 2021; Baym et al., 2016; Renz et al., 2025). However, the evolution of bacterial MDR involves many parallel instances of lineages acquiring (and losing) multiple coupled resistance features, challenging existing phylogenetic comparative methods (PCM) focussed on small numbers of coupled features (Boyko & Beaulieu, 2021; Pagel, 1994). Learning the evolutionary pathways by which bacteria acquire MDR, the features shaping these pathways, and future predictions of MDR emergence, remains an outstanding problem, despite existing and powerful studies across aligned theoretical and empirical topics (Baquero et al., 2021; Blanquart, 2019; Krieger et al., 2020; Lehtinen et al., 2019; Nichol et al., 2015; Palmer & Kishony, 2013).

Evolutionary accumulation modelling (EvAM), a branch of machine learning with its roots in cancer progression (Beerenwinkel et al., 2015), aims to infer co-evolutionary pathways involving multiple characters which can positively or negatively influence each others’ acquisition (Diaz-Uriarte & Herrera-Nieto, 2022; Diaz-Uriarte & Johnston, 2025). EvAM methods have been applied to drug resistance evolution in HIV (Beerenwinkel et al., 2005; Montazeri et al., 2015), *Mycobacterium tuberculosis* (Aga et al., 2024; Chen et al., 2026; Greenbury et al., 2020; Leandry et al., 2025; Moen & Johnston, 2023), and *Kp* (Aga et al., 2025), revealing patterns of convergent evolution and their dependence on geography and policy (Aga et al., 2025; Renz et al., 2025). In contrast with PCM, EvAM methods are not challenged by multiple features or their interactions, but often neglect feature loss and phylogenetic coupling between samples (Johnston & Diaz-Uriarte, 2025). HyperMk2 (Johnston et al., 2026), an EvAM variant of the established Mk model in evolutionary biology (Boyko & Beaulieu, 2021; Lewis, 2001; Pagel, 1994), has been recently introduced to address these shortcomings, supporting reversible and potentially simultaneous accumulation of multiple coupled binary characters in phylogenetically-embedded data. These methods allow the inference of transition networks as quantitative, comparative representations of evolutionary pathways, supporting predictions and analyse of influential features. Here, we use these new developments to learn the structure of evolutionary pathways to MDR across ESKAPEE pathogens, and thereby provide, and test, a predictive model for future MDR evolution in newly-observed infections.

## Results

### Evolutionary pathways across ESKAPEE pathogens

We obtained phenotypic data on drug resistance patterns in ESKAPEE pathogens from the CABBAGE database (Dickens et al., 2026) and estimated phylogenies using *mash* (Ondov et al., 2016) and genome data from NCBI (Sayers et al., 2024) (Fig. 1A; Supp. Fig. 1; see Methods). We then used HyperMk2 (allowing for reversible, potentially simultaneous acquisitions) and HyperHMM (Moen & Johnston, 2023) (using a picture of irreversible accumulation) to infer pathways of MDR evolution across ESKAPEE pathogen species, verifying consistency between the two approaches (Supp. Fig. 2). In agreement with previous work (Aga et al., 2025; Johnston, 2026), we found the orderings reported by HyperHMM to reflect the dynamics inferred under the more flexible picture of HyperMk2 (Supp. Fig. 2), meaning that an accurate representation of AMR evolutionary pathways (allowing for reversibility and multiple acquisitions) can be captured with the consistent visualisation and analysis supported by HyperHMM.

**Figure 1.**
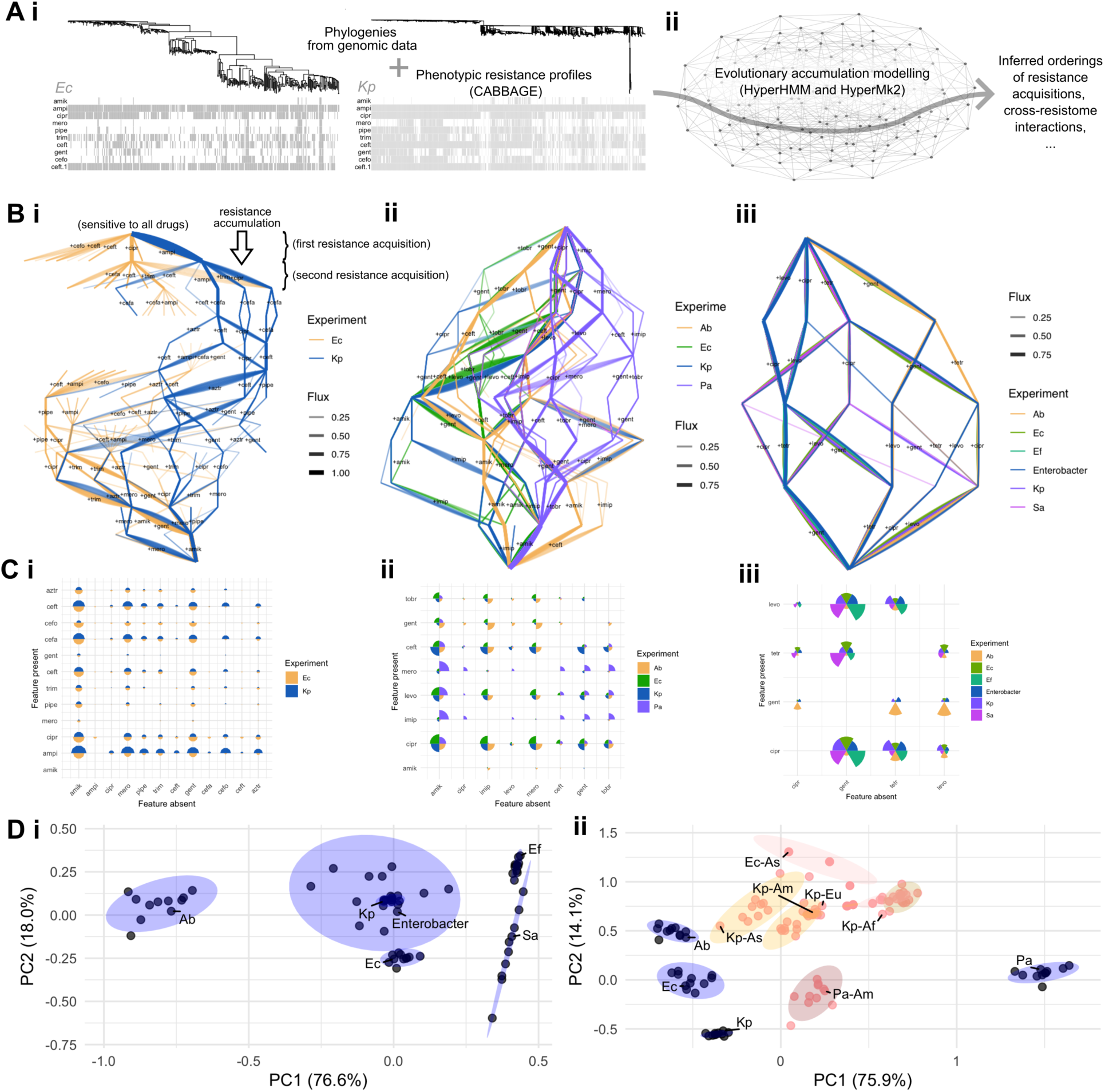
Shared evolutionary pathways to MDR across ESKAPEE pathogens. **(A)** Illustration of data and analysis pipeline (see Methods). Phenotypic resistome profiles from CABBAGE (Dickens et al., 2026), linked by phylogenies, are established for di]erent bacteria (here, *Ec* and *Kp*; see Supp. Fig. 1). Evolutionary accumulation modelling, a class of machine learning methods for inferring evolutionary dynamics through the space of resistomes (Renz et al., 2025), is used to learn the absolute and relative orderings of, and interactions between, resistance features. **(B)** Inferred transition networks from HyperHMM for drug resistance evolution across 10 drugs in *Ec* and *Kp* (i, Set 2-10); 8 drugs in 4 pathogens (ii, Set 4-8); and 4 drugs in 6 pathogens (iii, Set 6-4). Drugs are labelled by their first four characters (Supp. Table 1). From a pan-sensitive precursor state (top), resistances to di]erent drugs are accumulated down the network, with probability given by edge width. The range of edges corresponds to bootstrap resamples. **(B)** Relative ordering matrix of drug resistance. The size of a circle segment gives the probability that one feature (column) is absent when another (row) is present. **(C)** Principal component analyses (PCA) showing the main axes of variability in the HyperHMM ordering matrices in for Sets 6-4 (i) and 4-8 (ii). Di]erent points are bootstrap resamples of the data. Black points and blue ellipses correspond to global data; in (ii), the positions of ordering matrices subset by geographical region are also shown (*Af*, Africa; *Am*, Americas; *Eu*, Europe; *As*, Asia).

**Figure 2.**
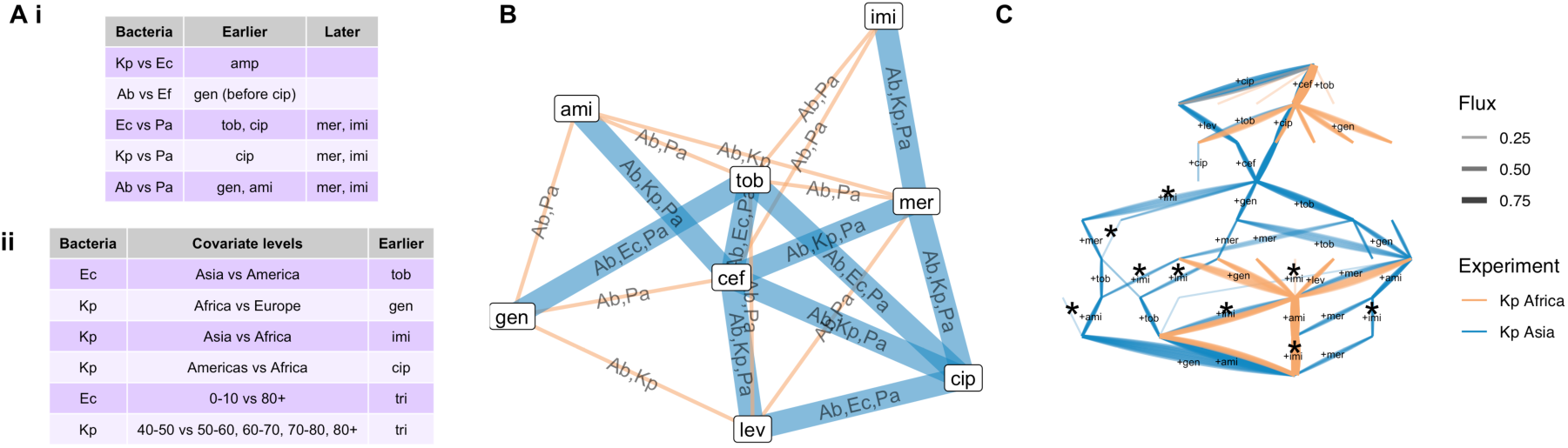
Drug interactions and covariates in global MDR evolutionary pathways. **(A)** (i) Strongest detected di]erences in inferred MDR pathways between species pairs; the drug(s) acquired earlier and later by X in the X vs Y pairing. (ii) Covariates shaping MDR evolutionary pathways. The species and covariate levels for which the strongest di]erences in drug acquisition ordering are inferred, and the drug(s) inferred to be acquired earlier by X in covariate value X vs Y. Drugs are labelled by their first three characters (Supp. Table 1). **(B)** Inferred interactions between drugs. Width and labels of edges gives number and identity of species for which a given interaction is identified (see also Supp. Fig. 4). **(C)** Example of transition network di]erences in inferred MDR dynamics in di]erent regions; imipenem resistance (* edges) has a statistically robust high probability of later acquisition in Africa (orange) than Asia (blue).

For each set of species we extracted the set of drugs for which most resistance information was available across the CABBAGE dataset, ensuring that diverse drug families were covered (Supp. Table 1). We focused on three subsets, labelled by number of species and drugs: Set 2-10 (10 drugs in *Ec* and *Kp*); Set 4-8 (8 drugs in *Ec*, *Kp*, *Ab* and *Pa*); and Set 6-4 (4 drugs in *Ec*, *Sa*, *Kp*, *Ab*, *Ef*, *Es*), and inferred pathways to MDR across species in each case. These pathways are reported as transition networks through an evolutionary space of phenotypic resistomes: patterns of resistance presence/absence across a palette of drugs (Fig. 1). Starting from a putative precursor state susceptible to all drugs, EvAM methods infer the likely ordering and relative timing of evolutionary steps by which new resistances are acquired (and lost) – annotated in Fig. 1Bi. The relative ordering of resistance acquisitions, and interactions between them (whether resistance to drug X increases or decreases the acquisition rate of resistance to drug Y) can then be established from the inferred pathways. Across diverse pathogens, resistance profiles will be shaped by different molecular mechanisms; our goal here is to identify and analyse similarities and differences at the phenotypic level (see Discussion). This approach also has the advantage of controlling for some possible biases in source data. Resistance profiles in AMR databases are likely enriched for unusually resistant bacteria, but as our approach considers pathways through the entire space of resistance patterns, inferred results will not be inappropriately weighted towards highly-resistance phenotypes.

We found that inferred MDR pathways are remarkably consistent across *Kp* and *Ec*, and to a large extent across these two and *Ab*, both in terms of the pathways through the space of resistance states (Fig. 1B), and the relative orderings of resistance acquisitions (Fig. 1C-D). Pathways in *Pa* and the other ESKAPEE species display some similar structures but with larger-scale differences, which can be individually characterised from the HyperHMM output (see Methods; Figs. 1C-D, 2Ai; Supp. Fig. 3). From Set 6-4, the relative orderings of gentamicin and ciprofloxacin are reversed in *Ab* and *Ef* (cipro first in *Ef*, genta first in *Ab*).

**Figure 3.**
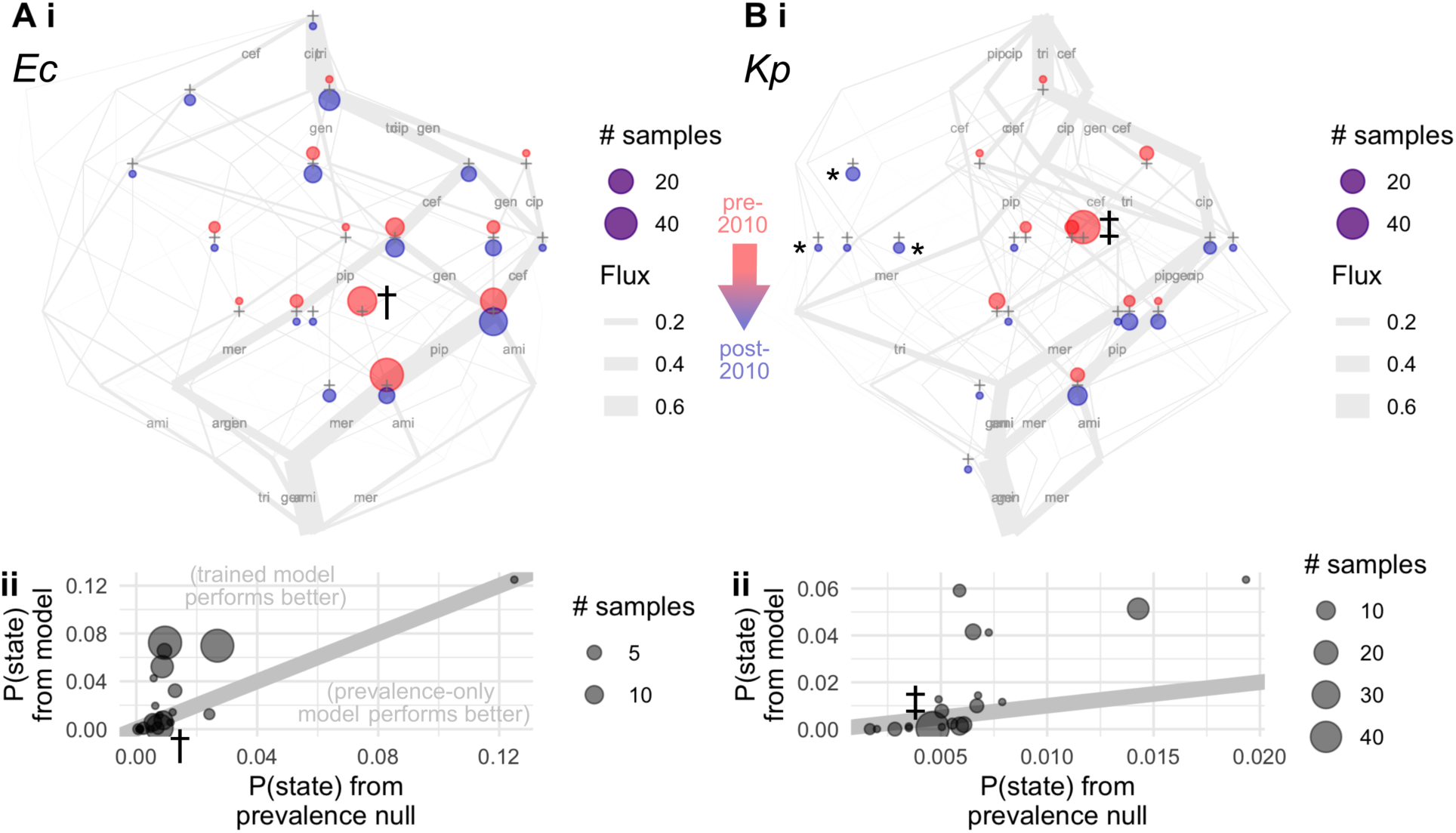
Predicted and observed MDR patterns in sub-Saharan *Ec* (A) and *Kp* (B) infections. (i) Grey, inferred transition networks (as in Fig. 1B) give the predicted evolution pathways from models on CABBAGE training data. Drugs are labelled by their first three characters (Supp. Table 1). Crosses give states observed in independent, newly-phenotyped bacterial isolates from Tanzania (see Methods). For each observed state, the size of upper red circles gives the number of observations in pre-2010 infections; the size of lower blue circles gives the number of observations in post-2010 infections. Specific states (see text): *, states with predicted probability < 10^-4^ from the trained model; †, gen-tri-cef-pip phenotype in pre-2010 *Ec*; ‡, gen-tri-cip-cef-pip phenotype in pre-2010 Kp. (ii) Probabilities of di]erent resistance states (with observation counts given by point size) from new phenotype data, under a null model considering only prevalence in the training data (horizontal axis; see Methods) and the trained HyperHMM model (vertical axis). † and ‡ correspond to the same states as in (i). Points above the grey line (y=x) correspond to observations with higher likelihood under the trained model.

From Set 4-8, relative to *Pa*, meropenem and imipenem resistance were acquired relatively late in *Ec*, *Kp* and *Ab*, ciprofloxacin resistance earlier in *Ec* and *Kp*, and gentamicin and amikacin resistance earlier in *Ab*. From Set 2-10, ampicillin resistance appears relatively early in *Kp* compared to *Ec* – a natural consequence of *Kp*’s general intrinsic resistance via SHV-type β-lactamases.

Across species and sets, HyperMk2 detected statistical support for interactions between drug resistances beyond a null model of independent character evolution (see Methods; Fig. 2B; Supp. Fig. 4). Not unexpectedly (though serving to sanity-check the inference), several of the drug interactions of highest inferred magnitude were within antimicrobial families and drug classes (tobramycin-gentamycin; meropenem-imipenem; levofloxacin-ciprofloxacin, respectively aminoglycosides, carbapenems, and fluoroquinolones; Supp. Fig. 4). However, we also detected interactions across pathogens between drugs from different families: (amikacin/tobramycin/levofloxacin)-ceftazidime and (meropenem/tobramycin)-ciprofloxacin. Each of these interactions reflects a departure from independent resistance acquisitions shaping MDR evolution: tob-cip interactions may, for example, be mediated by the presence of the *aac(6′)-Ib-cr* gene conferring resistance to both drugs (Jacoby et al., 2015), and shared resistance to other combinations can arise from the acquisition of mobile genetic elements containing multiple resistance genes (see Discussion).

### Covariates of evolutionary pathways

We next asked which external covariates influenced these inferred MDR pathways. In addition to bacterial species, the CABBAGE dataset has metadata including country of origin and demography of the patient from which a sample originates. Using bootstrapped replicates in HyperHMM (see Methods), we tested for differences in relative and absolute inferred orderings of MDR characters across different levels of these covariates (Figs. 1C, 2Aii; Supp. Fig. 5). We found statistical support for several influential covariates that refined the predictions of the evolutionary models for MDR pathways. For example, the relative ordering of imipenem resistance in *Kp* is later in Africa than other global regions, corresponding to previously inference of relatively late emergence of carbapenem resistance in Africa (Aga et al., 2025) (Fig. 2A,C). *Ec* acquires tobramycin resistance relatively earlier in Asia than Americas; *Kp* shows additional regional differences in resistance to gentamicin (earlier in Africa than Europe) and ciprofloxacin (earlier in the Americas than Africa). From the perspective of patient demographics, *Kp* acquires trimethoprim-sulfamethoxazole (bactrim) resistance relatively earlier in infections in 40-50 year-old patients than in 50-60, 60-70, and 70-80 year olds, with a likely contributing factor being common bactrim use in lower- and middle-income countries with younger populations.

### Testing predictions of MDR pathways in newly-phenotyped ESKAPEE infections over decades

We next asked whether EvAM models trained on existing data could predict MDR patterns in newly-observed ESKAPEE pathogens, and whether these predictions could be used prospectively to predict future evolutionary behaviour. To this end, we obtained phenotypic resistance profiles from several hundred isolates of *Kp* and *Ec* from clinical infections in Tanzanian hospitals between 2001 and 2018 (Supp. Fig. 6; see Methods). We compared the predictions from an EvAM model for each pathogen, trained on non-Tanzanian data, to the phenotypic profiles of these pathogen samples over time (Fig. 3).

Both pre-2010 and post-2010 observations fall overwhelmingly on the high-probability pathways predicted by the EvAM model (Fig. 3Ai, Bi), with several instances of observations compatible with a general flow of observation probability through the network from pre-2010 to post-2010 states. The likelihoods of observed states under the trained EvAM model are generally higher than under a null model based on resistance prevalence alone without feature interactions (Fig. 3Aii, Bii; see Methods). There were two exceptions. First, for a small number of observed states (marked * in Fig. 3) for which the model was not able to produce estimated probabilities due to an absence of training data. Once these points were removed, the trained models’ likelihoods associated with the new observations both exceeded the prevalence-only model (likelihood ratio 119.8 for *Ec*, 1.56 for *Kp*). Second, there are two notable states where the trained model predicted lower probabilities. The first is gen-tri-cip-cef-pip, a combination predicted with high probability in EvAM but even higher in the prevalence model, due to the absence of mer and ami in the training data. The second is gen-tri-cef-pip, a common resistance combination in pre-2010 observations that was not predicted with high probability from the trained model. This prevalence could be due to regionally or temporally constrained presence of plasmids conveying this resistance pattern that were less reflected in the CABBAGE training data (Daud & Nyombi, 2023). If these states were neglected, likelihood ratios increased further to 784.6 for *Ec*, 80.8 for *Kp*. Notably, this ability to capture unseen AMR evolutionary dynamics comes from phenotypic training data, without being informed by the genetic details of resistance mechanisms (which can also be used in more targetted EvAM analyses (Aga et al., 2025)).

## Discussion

We have used EvAM approaches to infer “roadmaps” of transition networks describing the evolutionary pathways to MDR across different drug sets and ESKAPEE pathogens. Several consistent principles appear throughout our analyses. Early resistance patterns correspond to common β-lactams and related families (penicillins, cephalosporins, and some fluoroquinolones). Intermediate resistance acquisitions include adjunct classes with members including piperacillin and bactrim, with resistance to last-line aminoglycosides and carbapenems appearing last. However, the EvAM analysis reveals a collection of higher-order effects which are not apparent from simple prevalences of these features (demonstrated quantitative by the AIC values favouring interaction terms in the analysis). Positive interactions between resistances in different antibiotic classes are identified across pathogens, and influence of geographical and demographic covariates on MDR evolutionary dynamics, are statistically supported despite the heterogeneous source data. Predictions from these higher-order trained models are supported by new phenotypes from historical data. Particularly as covariates like drug use statistics can sometimes display limited predictive power for AMR evolution (Aga et al., 2025), these results support the possible contributions of this approach in forecasting future patterns of MDR in different pathogens (Blanquart, 2019; Olesen et al., 2018).

The fundamental modelling technology in EvAM is a Markov model, which also underpins modern AI workflows. It is of interest to note a conceptual similarity with exciting recent AI work on, for example, the natural history of human disease (Shmatko et al., 2025). There, a (much more sophisticated) approach tokenises disease-aligned states and predicts token sequences over time. Our approach, particularly in its predictive application, has a similar philosophy of predicting the next “token” drug resistance from a given state. As with other machine learning approaches, our analysis here will improve as training data expands, especially with respect to covariate-driven differences in MDR evolutionary pathways. One example would be analysing resistome patterns inferred from genomic data; here, for a direct experimental connection, we have focussed on patterns directly derived from laboratory observations, but including genome-based estimates reflects another possible line of enquiry (Aga et al., 2025; Davis et al., 2016; Palmer & Kishony, 2013; Ren et al., 2022; Su et al., 2019). While our focus here is on a broad, comparative analysis of phenotypic evolutionary patterns, a future genomic perspective will support more detailed mechanistic analysis of resistance evolution, including distinguishing specific molecular mechanisms, and plasmid-driven resistance from other types (Darby et al., 2023; Holmes et al., 2016).

While large-scale datasets are supporting powerful analyses of AMR across bacteria and regions (Holt et al., 2015; Munk et al., 2022; Wyres et al., 2020), it has been argued that “knowledge regarding ‘what happened’ has precluded a deeper understanding of ‘how’ evolution has proceeded” (Baquero et al., 2021). Here we have attempted to reconcile these two pictures, using broad and deep data to learn the transition networks describing MDR evolutionary pathways (Boyko et al., 2026; Renz et al., 2025). In treating these networks as the central, quantitative unit of analysis, we have shown that universal behaviours, species-specific differences, covariates, and predictions can all emerge from an evolutionary picture unifying diverse individual observations (Aga et al., 2025).

## Methods

### Global phenotype data

We use the CABBAGE dataset (Dickens et al., 2026) for phenotypic drug resistance patterns across different bacterial samples. CABBAGE contains experimentally measured resistant/susceptible data for bacteria along with accessions to the corresponding genomes. For a given comparison experiment, we first choose the set of bacterial species under investigation. We then rank the drugs in CABBAGE by number of records in this set. For the top-ranked *n* drugs, we identify the collection of records in CABBAGE that have complete resistant/susceptible information across the set of bacteria for each of the *n* drugs.

### Phylogenetic relationships

To allow our pipeline to be reproducible with a single machine, we used an approach that limits both the amount of processor time and storage space required. We retrieved the corresponding genome accessions from NCBI (Sayers et al., 2024), in batches of 400 (roughly 2Gb) at a time. We used *mash* (Ondov et al., 2016) to “sketch” each downloaded genome, storing sketches rather than full genome information. When all sketches for a given dataset had been computed, we used *mash* to compute a matrix of summary distances between each. This distance matrix was then used in a simple neighbour-joining algorithm in *ape* (Paradis & Schliep, 2019) in R (R Core Team, 2022) to estimate a coarse-grained phylogeny.

### Learning evolutionary pathways and interactions

The matrix of resistant/susceptible markers and this estimated phylogeny were then used as input either to HyperMk2 (Johnston et al., 2026) or HyperHMM (Moen & Johnston, 2023), approaches for evolutionary accumulation modelling (Diaz-Uriarte & Herrera-Nieto, 2022; Diaz-Uriarte & Johnston, 2025; Renz et al., 2025). Briefly, HyperMk2 estimates a minimum-evolution state space for a given set of observations and infers a transition matrix between these states most compatible with data. The fitted model was compared to a null model of independent character evolution and interactions are estimated through comparison with pairwise Mk models (Johnston & Diaz-Uriarte, 2025). Here, we report ΔAIC_model_ between this null model and the fitted model. We then report ΔAIC_i,j_ between a model where i and j are codependent and a null model where they evolve independently (Boyko & Beaulieu, 2021; Grundler, 2025; Johnston et al., 2026). We report cases where ΔAIC_i,j_ > ΔAIC_model_/L, where L is the number of characters, to identify the pairwise interactions that most strongly contribute to the coupled model’s strength. HyperHMM estimates the weights on a hypercubic network of irreversible transitions most compatible with observations and an ancestral state reconstruction based on irreversible, rare transitions (Moen & Johnston, 2023).

### Transition network differences

We analysed both relative and absolute orderings of feature acquisitions in an inferred transition network. Relative ordering 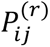 gives the probability that feature *i* is acquired before specific feature *j*. Absolute ordering 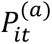 gives the probability that feature *i* is acquired at step *t* in a pathway. For the PCA plots comparing network dynamics, 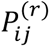 was concatenated to a vector for each bootstrap sample for each case. Differences in the relative and absolute inferred orderings of character acquisitions can be tested across bootstrap resamples. We report two classes of difference: first, cases of ordinal time *t* where 95% or more of the bootstrapped distributions support one species acquiring the drug before *t* with ≥75% probability and the other species acquiring the drug after *t* with ≥75% probability. Second, cases of pairs of drugs X and Y where 95% or more of the bootstrapped distributions support X being acquired before Y in one species and Y before X in the other.

### New bacterial phenotypes

Phenotypes from *Kp* and *Ec* isolates were derived from a collection of studies performed in Tanzania and Zanzibar over the last three decades. These comprise bacterial isolates from (a) blood samples from pediatric patients with bloodstream infections in Dar-es-Salaam, Tanzania in 2001-2002 (Blomberg et al., 2005); (b) blood samples from adult patients with bloodstream infections in Dar-es-Salaam, Tanzania in 2017-2018 (Moyo et al., 2020); (c) fecal samples from the patients in (b) (Kibwana et al., 2022) ; and (d) blood samples from adult and pediatric patients with bloodstream infections in Mnazi Mmoja hospital, Zanzibar in 2015-2016 (Onken et al., 2024) . Susceptibilities against a panel of 12 antimicrobial agents (Supp. Fig. 6) were tested by the disk diffusion method according to (a) NCCLS guidelines (National Committee for Clinical Laboratory Standards, 1997) (b, c) guidelines and breakpoints of the Clinical Laboratory Standards Institute (CLSI) (Clinical and Laboratory Standards Institute, 2015); (d) the EUCAST guidelines (EUCAST, 2022), with further details in the individual publications describing the sample collection. For forecasting experiments we used the 7 drugs for which most complete records appeared in the CABBAGE training data: gentamicin, trimethoprim-sulfamethoxazole, ciprofloxacin, ceftazidime, piperacillin-tazobactam, meropenem, and amikacin.

### Prevalence-only ordering model

The prevalence-only null model is derived from a weighted Plackett-Luce model for sequential accumulation with random stopping time. The weights for the Plackett-Luce process are taken as the prevalence probabilities of each feature in the training data. The likelihood of an observation with k features acquired S = {S_1_, … S_k_} is then L = P(k) P(first k features of the weighted Plackett-Luce process are the elements of S), where P(k) is the probability that a k is chosen as the random stopping time in the process (which is taken to be a uniform integer between 0 and 7, the total number of features, to match the HyperHMM emission model), and the feature probability is computed over permutations by dynamic programming.

## Code and data availability

All newly-compiled phenotype data, and code for the analysis, is freely available at https://github.com/StochasticBiology/EvAM-ESKAPEE. In addition to the packages above, the code uses R (R Core Team, 2022) with packages *arrow* (Richardson et al., 2023), *readxl* (Wickham & Bryan, 2019) for data entry, *dplyr* (Wickham et al., 2023), *tidyr* (Wickham et al., 2020) for data curation, *igraph* (Csardi & Nepusz, 2006), *stringr* (Wickham, 2019) for processing, and *ggplot2* (Wickham, 2011), *ggrepel* (Slowikowski, 2021), *ggpubr* (Kassambara, 2020), *ggraph* (Pedersen, 2020), *ComplexUpset* (Lex et al., 2014) for visualisation.

## Acknowledgements

This project has received funding from the European Research Council (ERC) under the European Union’s Horizon 2020 Research and Innovation Programme [Grant agreement No. 805046 (EvoConBiO) to I.G.J.]. This project was supported by the Trond Mohn Foundation [project HyperEvol under grant agreement No. TMS2021TMT09] to I.G.J., through the Centre for Antimicrobial Resistance in Western Norway (CAMRIA) [TMS2020TMT11] to N.L. This project has received support from ERA-Net: JPI-AMR STRESST project (NFR333432) to N.L.

---

**Supplementary Figure 1.**
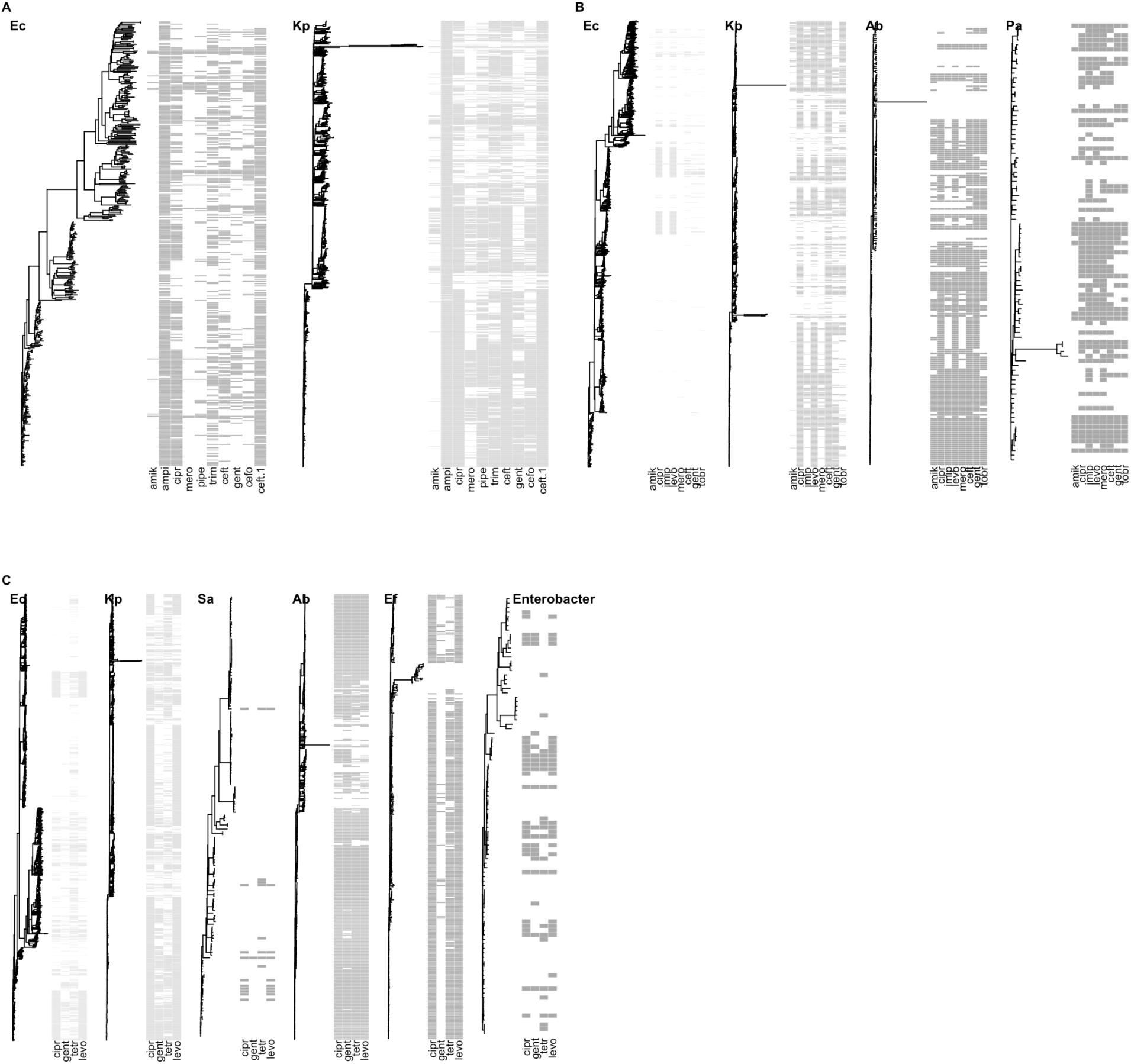
Estimated phylogenies and phenotypic resistance profiles for the CABBAGE database subsets in our study. (A) Set 2-10; (B) Set 4-8; (C) Set 6-4. For each species, the estimated phylogeny and presence/absence (grey/white pixels) patterns for drug resistances are shown. Drugs are labelled by their first four characters (Supp. Table 1); “ceft” and “ceft.1” correspond to ceftazidime, ceftriaxone.

**Supplementary Figure 2.**
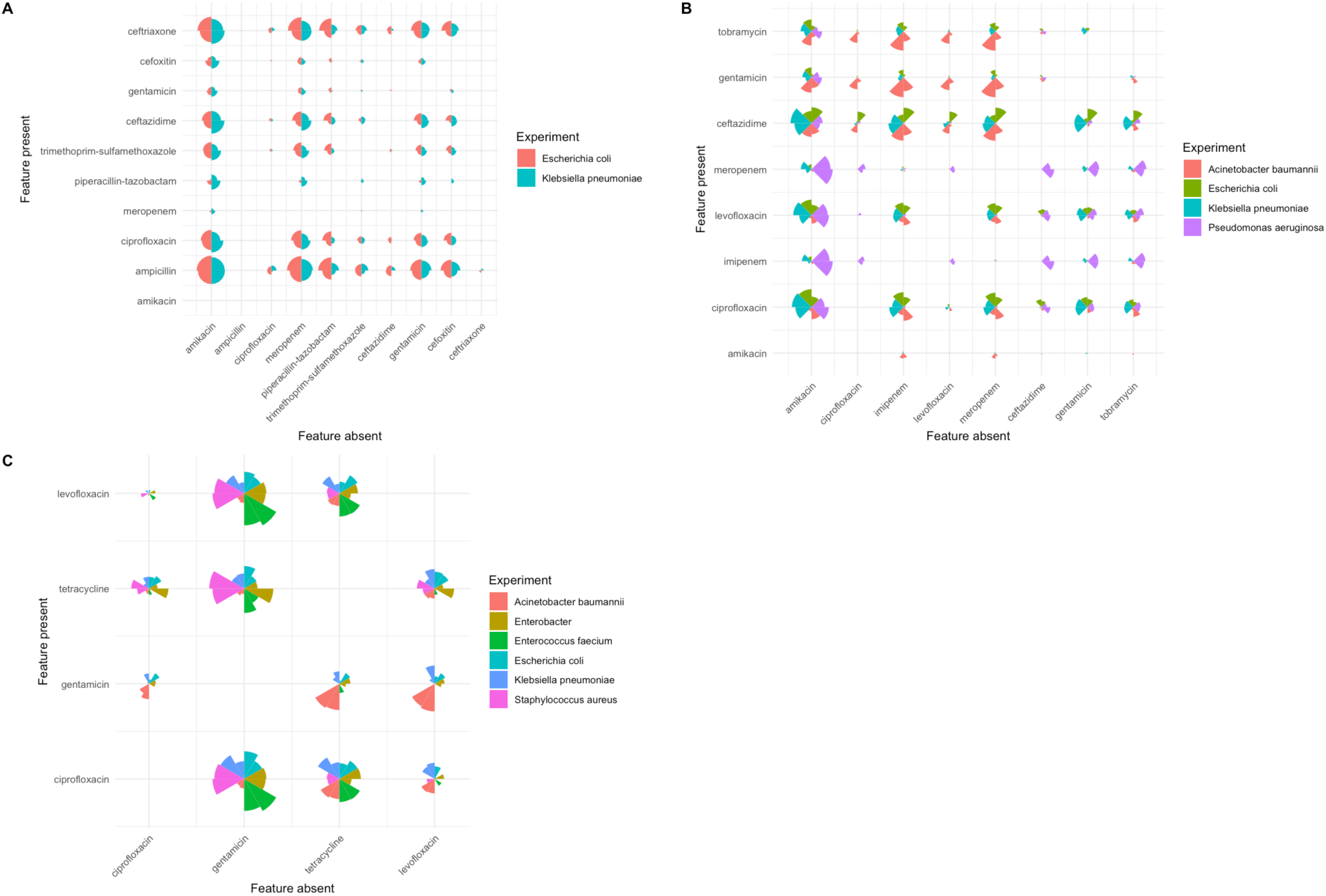
MDR pathways inferred with HyperMk2 and HyperHMM. In these “relative ordering matrices” (see Methods), the radius of each circle segment gives the probability that resistance to a given drug (column) is absent when resistance to another drug (row) is present. Each matrix has two segments of the same colour for each species; the clockwise segment is from HyperHMM and the anticlockwise segment is from HyperMk2. Both inference methods agree in the relative orderings of resistance acquisitions across pathogens. Comparisons are for **(A)** dataset 2-10; **(B)** dataset 4-8; **(C)** dataset 6-4.

**Supplementary Figure 3.**
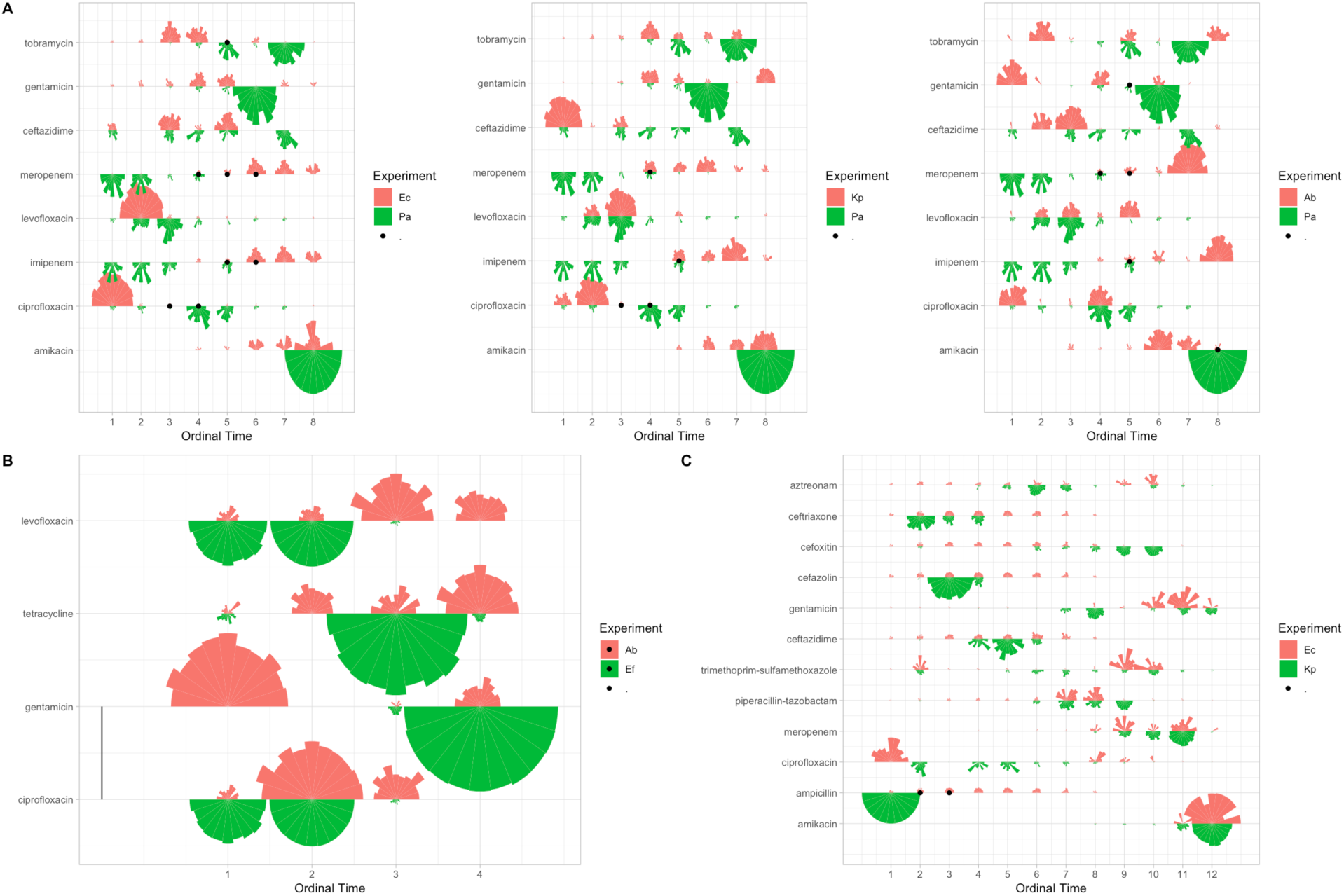
Differences in MDR evolutionary pathways between pairs of ESKAPEE pathogens. **(A)** Differences from dataset 4-8; **(B)** differences from dataset 6-4; **(C)** differences from dataset 2-10. Experiment The radius of each circle segment gives the probability that resistance to a given drug (row) is acquired at a given ordering in an evolutionary pathway (column). Different circle segments give probabilities from different bootstrap resamples, different colours give different bacterial species. A black dot at ordinal time t shows cases where 95% or more of the bootstrapped distributions support one species acquiring the drug before t with ≥75% probability and the other species acquiring the drug after t with ≥75% probability. Vertical bars give pairs of drugs X and Y where 95% or more of the bootstrapped distributions support X being acquired before Y in one species and Y before X in the other.

**Supplementary Figure 4.**
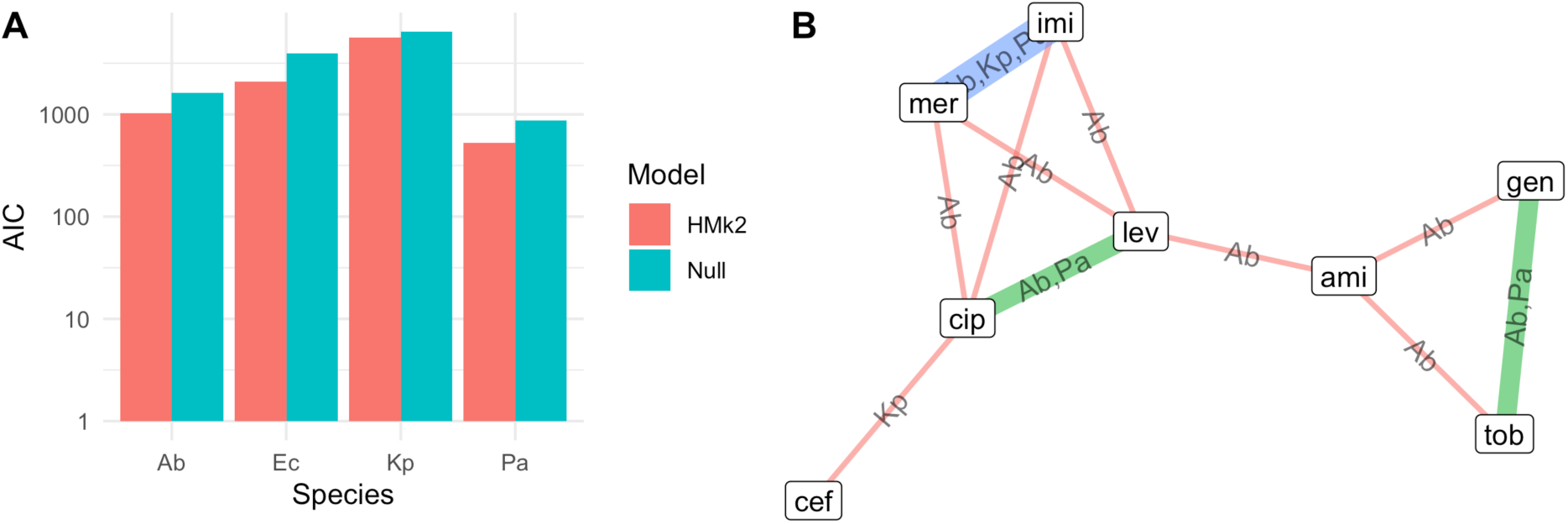
Inferred interactions between drug resistances in Set 4-8. **(A)** AIC values for the HyperMk2 model allowing for interactions (HMk2) and a null model of independent character evolution (Null). **(B)** Detected interactions with ΔAIC_i,j_ > ΔAIC_model_/L. The width of an edge gives the number of species in which the interaction was detected. Drugs are labelled by their first three characters (Supp. Table 1).

**Supplementary Figure 5.**
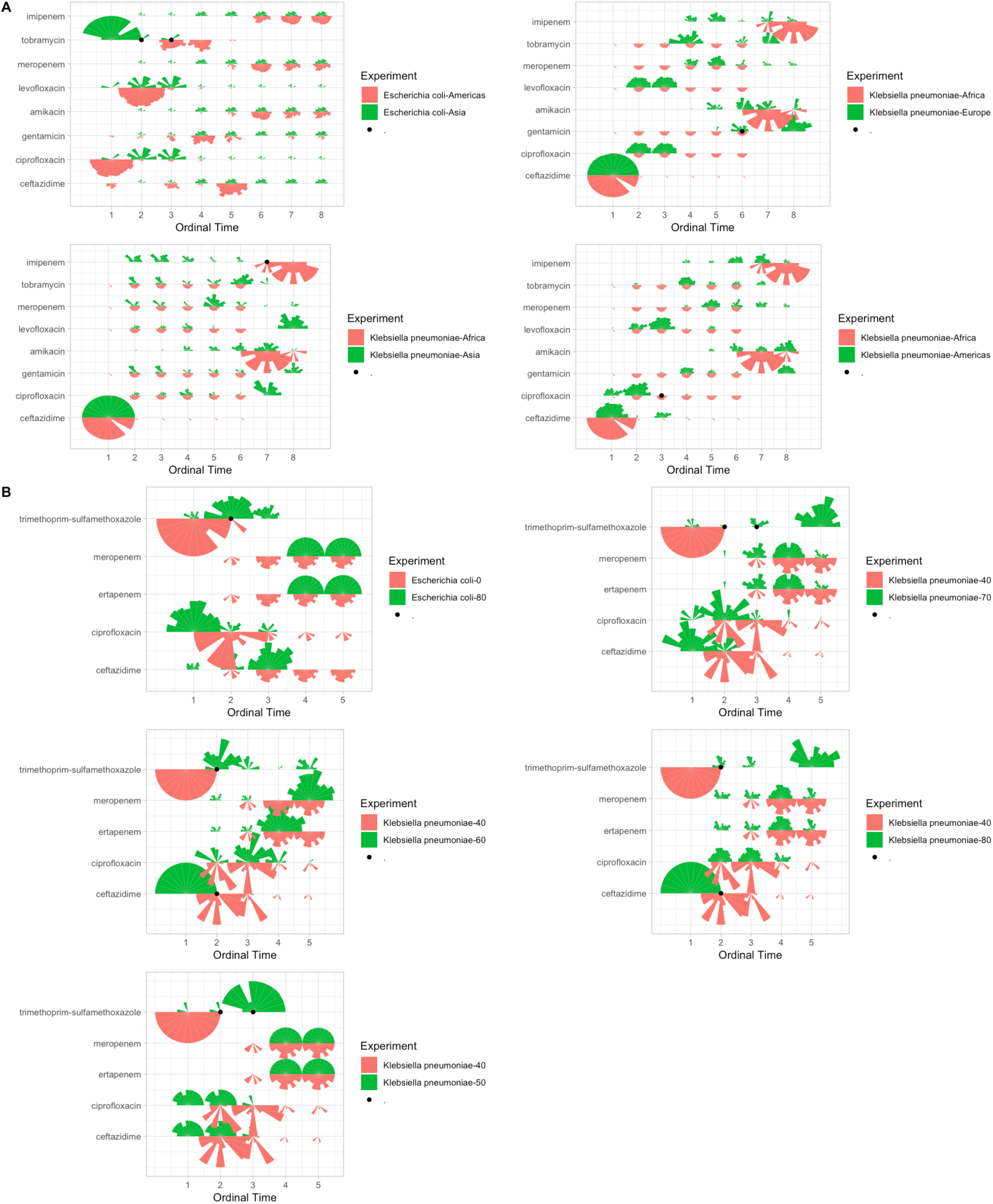
Covariate levels shaping MDR pathway evolution. **(A)** Continent of origin; **(B)** age bracket (in decades) of host patient. The radius of each circle segment gives the probability that resistance to a given drug (row) is acquired at a given ordering in an evolutionary pathway (column). Different circle segments give probabilities from different bootstrap resamples, different colours give different bacterial species. A black dot at ordinal time t shows cases where 95% or more of the bootstrapped distributions support one species acquiring the drug before t with ≥75% probability and the other species acquiring the drug after t with ≥75% probability.

**Supplementary Figure 6.**
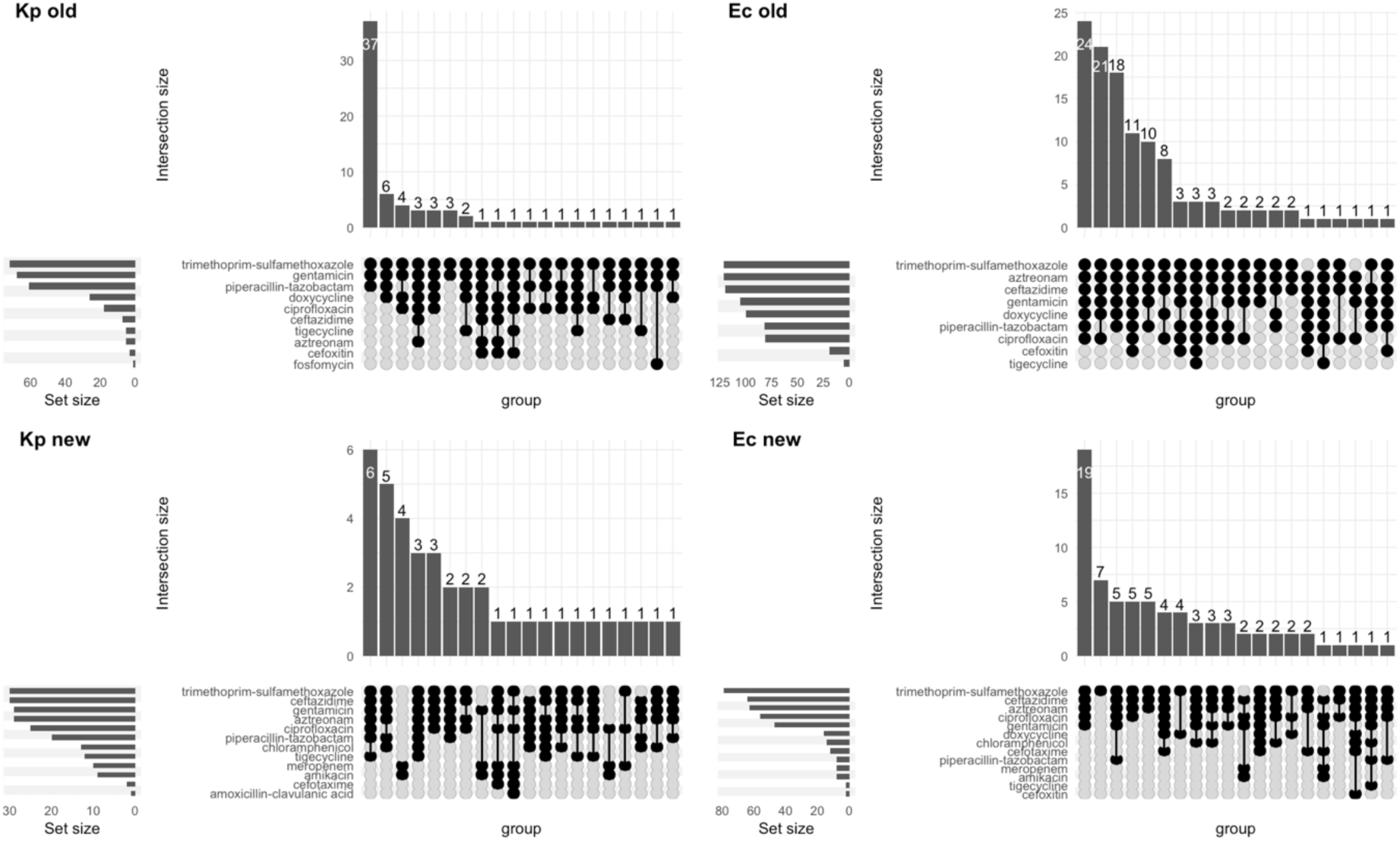
Most-common resistance patterns in the newly-phenotyped ESKAPEE samples. The four classes are labelled by species and “old” (pre-2010) vs “new” (post-2010). In each case, the UpSet plots (Ahlmann-Eltze, 2025) show the 20 most common patterns of drug resistance (black circles denote resistance) with total count for each pattern (vertical bars) and total independent count of each drug resistance in the sample (horizontal bars).

**Supplementary Table 1.**
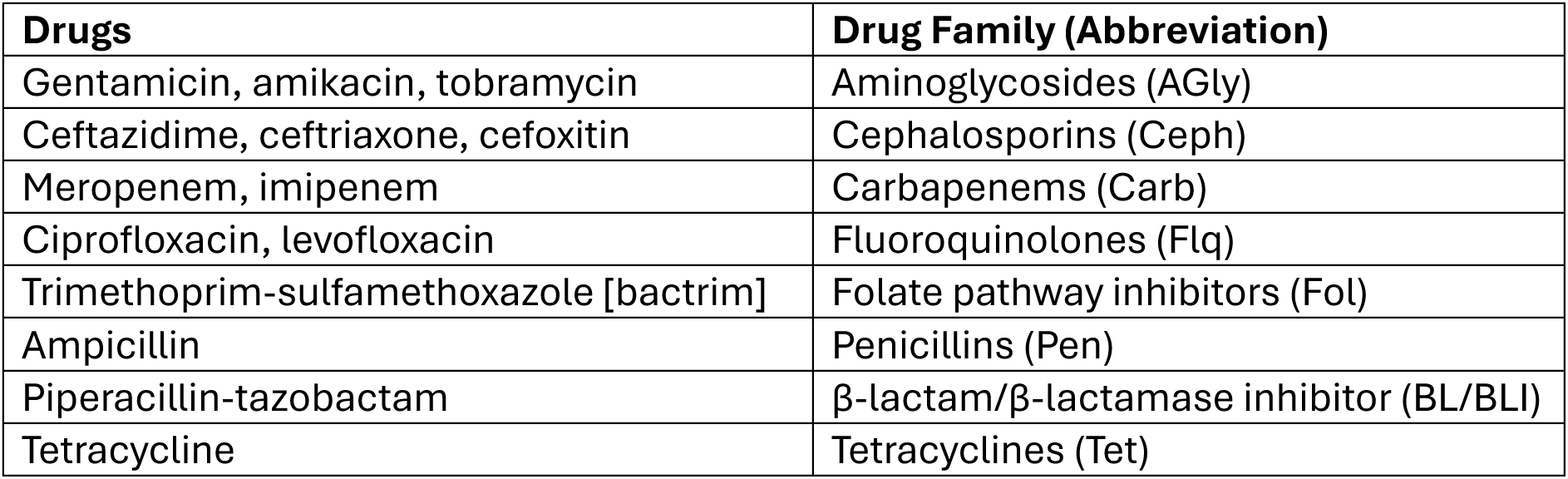
Specific drugs mentioned in this research, and their corresponding families.

